# The tetraspanin disc proteins, peripherin-2 and ROM1, facilitate CNG channel localization to the rod outer segment

**DOI:** 10.1101/2025.09.30.679618

**Authors:** Molly T. Thorson, Stephanie E. Wei, Junseo Park, Jorge Y. Martínez-Márquez, David G. Ball, Jason R. Willer, William J. Spencer, Jillian N. Pearring

## Abstract

The light-responsive outer segment of rod photoreceptors is composed of two distinct membrane subdomains: discs and the plasma membrane. We investigate how the disc protein peripherin-2 is engaged in CNG channel delivery to the outer segment. Instead of forming outer segments, peripherin-2 knockout (*Rds*^*-/-*^) photoreceptors release ciliary ectosomes, in which CNG channel levels are markedly reduced relative to other outer segment proteins. This is intriguing as downregulation of the CNG channel is not a general feature of degenerative mouse models with dysmorphic outer segments. Overexpression of the β1-subunit of CNG in *Rds*^*-/-*^ rods reveals that the majority is trapped in intracellular membranes, but is restored to the outer segment by co-expressing peripherin-2. We test peripherin-2 chimeras containing either the N-terminus, tetraspanin core, or C-terminus and find that the tetraspanin domain is sufficient to localize CNGβ1 to the outer segment. We further show that the membrane remodeling function of the tetraspanin domain facilitates this process by the redundant action of the tetraspanin domain from ROM1 and the reemergence of the endogenous CNG channel in aged *Rds*^*-/-*^ rods that have produced ciliary membrane protrusions.

## INTRODUCTION

Human vision begins when photons of light initiate a biochemical signal that elicits an electrical response within the outer segment compartment of photoreceptor cells. The outer segment is a cylindrical organelle filled with hundreds of flattened membranes called discs, which house visual transduction machinery and structural proteins. This organelle has the singular function of generating the light response and therefore does not contain normal protein turnover machinery. To replenish membrane proteins, discs must be constantly renewed through the addition of new discs at the base of the outer segment and subsequent phagocytosis of old discs at the distal tip by the overlying retinal pigment epithelium (RPE) (1). The entire outer segment is renewed over an approximately two-week time span (2), placing a high demand for membrane trafficking of both signaling and structural proteins into the outer segment (3, 4). Defects in disc renewal or membrane trafficking underlie many forms of inherited retinal degeneration that cause human blindness.

In rod photoreceptors, disc membranes are physically separate from the plasma membrane surrounding them. Protein composition of these two membrane subdomains is distinct. Disc membranes house proteins that initiate the visual response, as well as the structural proteins, peripherin-2 and ROM1, which are localized to the hairpin curve of the disc rim. Proteins that modulate the electrical properties of the cell are restricted to the plasma membrane (5-7). It has been suggested that discs are stabilized laterally to the plasma membrane through interactions with disc rim proteins. The prime candidates for this interaction are peripherin-2 and β1-subunit of the CNG channel (CNGβ1) based on evidence that peripherin-2 has been detected following immunoprecipitation of CNG channel subunits (8). Membrane proteins must therefore be sorted into these membrane subdomains based upon their function. To better understand how membrane proteins populate the plasma membrane, we examined mechanisms of CNG channel delivery to the outer segment.

The CNG channel is a heterotetramer that consists of three α1- and one β1-subunit in rod photoreceptors (9, 10). While the α1-subunit (CNGα1) can form functional homotetramers in the absence of CNGβ1 (11), the reverse is not true (12). The 3:1 stoichiometry of the channel subunits is critical for proper localization and function of the CNG channel in the outer segment. In fact, the channel subunits have been shown to assemble early in biosynthesis and traffic as a fully formed channel to the plasma membrane of the outer segment (13). Previous studies found that in the absence of CNGβ1, CNGα1 is severely downregulated (14, 15). When CNGα1 is overexpressed in CNGβ1-deficient rods, it does not target specifically to the plasma membrane of the outer segment, and instead, localizes indiscriminately throughout the plasma membrane of the cell, suggesting that CNGα1 does not contain targeting information (13). It was shown that the outer segment targeting information for the CNG channel is contained within the β1 subunit (13).

To get to the outer segment, membrane proteins utilize two trafficking routes that diverge shortly after biosynthesis in the ER. The conventional trafficking route utilizes vesicular transport through the Golgi prior to reaching the outer segment. Membrane proteins that utilize the unconventional trafficking route bypass the Golgi, relying on vesicular transport from the ER to reach the outer segment. Proteins such as rhodopsin and the CNG channel have been identified to traffic conventionally, while the disc rim proteins peripherin-2 and ABCA4 use the unconventional trafficking route to reach the outer segment (13, 16-19). In mice, there is a small proportion of peripherin-2 that traffics through the conventional pathway, which was shown to require its binding partner and homolog, ROM1 (20). The biological importance of segregating disc rim proteins into each trafficking pathway remains unknown.

In the peripherin-2 knockout mouse (aka retinal degeneration slow, *Rds*), photoreceptor outer segment proteins and membrane material are not incorporated into discs, but instead are released in extracellular vesicles called ectosomes (21). Curiously, it was reported that the CNGβ1 subunit was virtually undetectable in peripherin-2 knockout retinas, suggesting that the CNG channel may require peripherin-2 for delivery to the outer segment (22, 23). Supporting this possibility, a previous study using bimolecular fluorescence complementation assays in transgenic *Xenopus* rods showed that overexpressed peripherin-2 and CNGβ1 successfully complement in the inner segment where protein synthesis occurs (24). In this study, we investigated whether peripherin-2 facilitates CNG channel localization to the outer segment by performing a comprehensive analysis of channel expression and localization in *Rds*^*-/-*^ retinas. We found that CNGβ1 is detectable in ectosomes released from the photoreceptor cilia of young *Rds*^*-/-*^ mice, though at much lower levels than other disc-specific proteins. Interestingly, we find progressive accumulation of CNGβ1 in the outer segment region of *Rds*^*-/-*^ retinas at a timepoint when photoreceptors begin to elaborate membrane whorls from their cilia. Since CNGβ1 contains the outer segment targeting signal for the channel, we went on to investigate peripherin-2’s role in CNGβ1, and therefore channel, delivery. We find that trafficking of the CNG channel does not require peripherin-2/ROM1 complexes that utilize the conventional pathways through the Golgi. Further, we found that the membrane-spanning tetraspanin region of peripherin-2 is sufficient to rescue outer segment localization of the channel. We also discovered that the tetraspanin core of ROM1 can facilitate CNGβ1 localization, congruent with these tetraspanin proteins having redundant functions.

## RESULTS

### In aged *Rds*^*-/-*^ rods, CNG channel localization is restored to lamellar membrane outgrowths

We characterized CNG channel subunit expression and localization in a side-by-side comparison between WT, *Rds*^*+/-*^, and *Rds*^*-/-*^ retinas at P21 before the onset of photoreceptor degeneration. We performed immunofluorescence staining on WT and *Rds*^*-/-*^ retinal sections (*Rds*^*+/-*^ retinal sections found in Supplemental Figure 1A) and found CNGα1 and ROM1 localized to the outer segment region (Figure 1A). To detect CNGβ1, we used an anti-GARP antibody that recognizes the N-terminal GARP domain of CNGβ1. This antibody also recognizes two soluble GARP1 and GARP2 proteins that are expressed from the *Cngb1* gene as a result of alternative mRNA splicing (25). GARP proteins have normal outer segment localization in WT and *Rds*^*+/-*^ retinas, while no fluorescence was observed in the outer segment region of *Rds*^*-/-*^ retinal sections at P21 (Figure 1A). We corroborated the apparent absence of CNGβ1 in *Rds*^*-/-*^ retinal sections using a CNGβ1-specific antibody (Supplemental Figure 2). qRT-PCR analysis from retinal lysates revealed that *Cnga1, Cngb1 Rom1, and Abca4* mRNA expression were reduced by ∼50% in *Rds*^*-/-*^ compared to WT or *Rds*^*+/-*^ retinas (Figure 1B). This suggests a global downregulation of outer segment components in *Rds*^*-/-*^ retinas. We then quantified protein levels of CNGβ1, CNGα1, ROM1, and ABCA4 from serial dilutions of retinal membrane fractions by Western blotting. We found that protein levels of CNGα1, CNGβ1 and ROM1 were drastically reduced compared to ABCA4, a protein previously shown to be present in *Rds*^*-/-*^ ectosomes (Figure 1C,D)(26). We were unable to detect either α1 or β1 subunits of the CNG channel by Western blot analysis of *Rds*^*-/-*^ retina membrane fractions. To determine if there is even a small amount of CNG channel delivered to the cilium (and therefore in ectosomes), we purified ectosomes from multiple P21 *Rds*^*-/-*^ retinas and analyzed this highly concentrated preparation by Western blotting. By this method, we were able to detect both CNGα1 and CNGβ1 in ectosomes (Figure 1E). Likewise, a previous study conducted mass spec analysis of purified *Rds*^*-/-*^ ectosomes and detected CNGα1 along with virtually all other outer segment proteins. However, this previous study did not include an analysis of CNGβ1 (26). We revisited this proteomic data and confirmed the presence of CNGβ1 in the *Rds*^*-/-*^ ectosome proteome (Supplemental Table 1). Together, our data show that compared to other outer segment membrane proteins, such as ABCA4, the CNG channel is severely downregulated in *Rds*^*-/-*^ retinas. We hypothesized that the inability to form an outer segment in the absence of peripherin-2 specifically influences either CNG channel stability or delivery.

**Figure 1.**
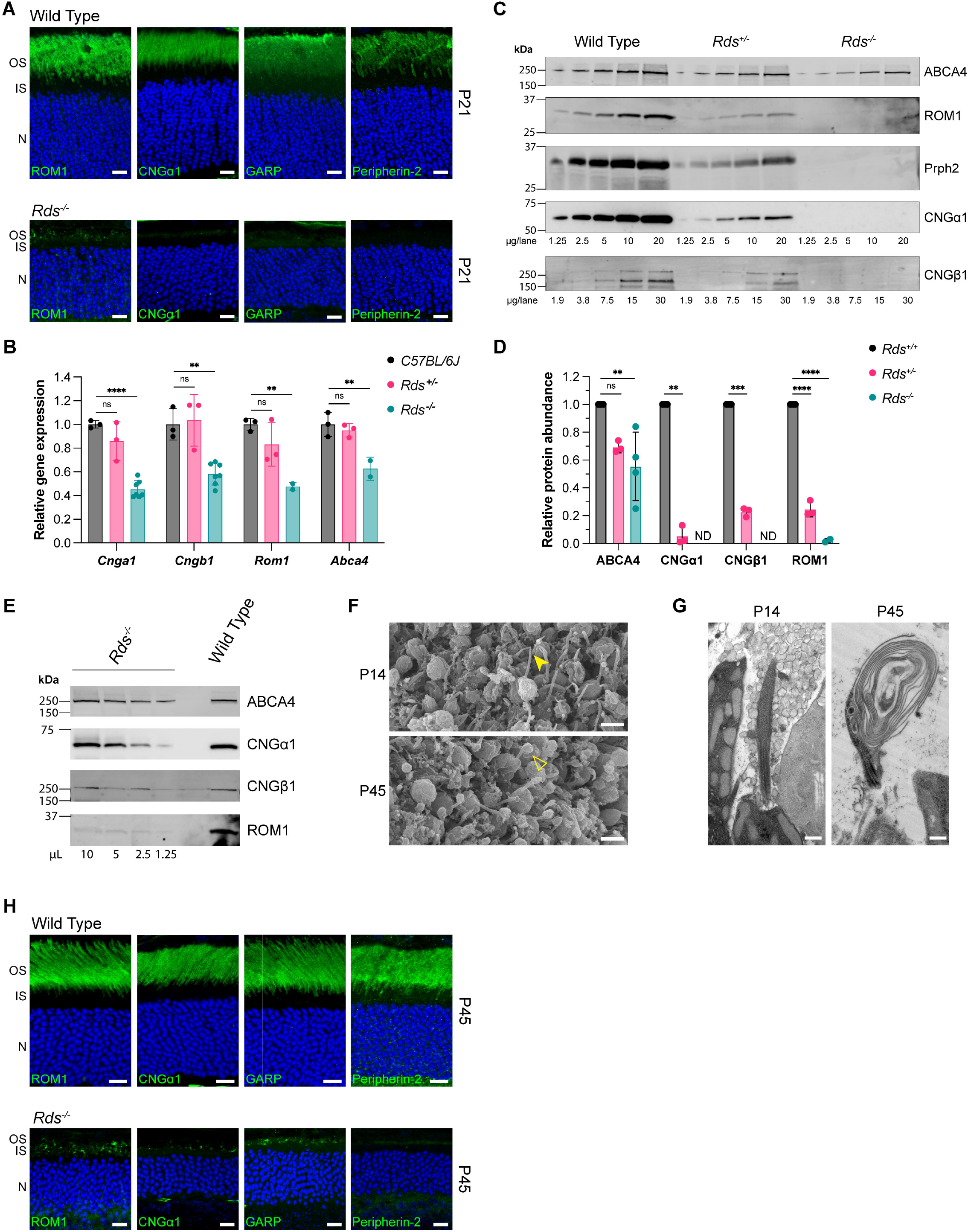
Characterization of CNG channel subunit localization in *Rds*^*-/-*^ retinas. **A**. Retinal sections from WT and *Rds*^*-/-*^ mice at P21 immunostained for ROM1, CNGα1, GARP proteins, and peripherin-2. Nuclei counterstained in blue. Scale Bar, 10 µm. Here and throughout the manuscript: outer segment (OS), inner segment (IS), and nucleus (N). **B**. *Cnga1, Cngb1, Rom1*, and *Abca4* mRNA levels in P21 WT, *Rds*^*+/-*^, and *Rds*^*-/-*^ retinal lysates were determined by qRT-PCR analysis. One-way ANOVA performed. pValues for *Cnga1* ****, p<0.0001; ns, p=0.152; *Cngb1* **, p=0.0021; ns, p=0.923; *Rom1* **, p=0.0092; ns, p=0.246; *Abca4* **, p=0.0087, ns=0.718. **C**. Representative Western blots show serial dilutions of WT, *Rds*^*+/-*^, and *Rds*^*-/-*^ membrane fractions for ABCA4, ROM1, Prph2, CNGα1, and CNGβ1. **D**. Relative protein expression is plotted in the bar graph. One-way ANOVA performed. pValues for ABCA4 ns, p=0.0525; **, p=0.0059 and ROM1 ****, p<0.0001. Welch’s t-test performed. pValues for CNGα1 **, p=0.0016; and CNGβ1 ***, p=0.0006. **E**. Representative Western blots show serial dilutions of an *Rds*^*-/-*^ ectosome fraction for ABCA4, CNGβ1, CNGα1, and ROM1 compared to a WT membrane fraction at a single concentration. **F**. SEM images of the apical surface of *Rds*^*-/-*^ retinas at P14 and P45. Scale Bar, 2 µm. Closed yellow arrowhead indicates photoreceptor cilia in young *Rds*^*-/-*^. Open yellow arrowhead indicates membrane mass protruding from photoreceptor cilia in aged *Rds*^*-/-*^. **G**. TEM images of *Rds*^*-/-*^ retinal cross-sections at P14 and P45. Scale bar 0.5 µm. **H**. Retinal sections from WT and *Rds*^*-/-*^ mice at P45 immunostained for ROM1, CNGα1, GARP proteins, and peripherin-2. Nuclei counterstained in blue. Scale Bar, 10 µm.

In the original publications characterizing the ultrastructure of *Rds*^*-/-*^ photoreceptors at P21, it was reported that a laminar membrane protrusion was occasionally seen emanating from the photoreceptor cilium (27-29). We performed scanning electron microscopy (SEM) from *Rds*^*-/-*^ retinas at P14 and P45 to assess the global appearance of photoreceptor cilia over time. By en face imaging of the apical surface of photoreceptor cells in young *Rds*^*-/-*^ retinas, we observed photoreceptor cilia surrounded by extracellular vesicles (Figure 1F). In contrast, many photoreceptor cilia of aged *Rds*^*-/-*^ retinas develop an enlarged morphology (Figure 1F). We performed TEM of *Rds*^*-/-*^ retinas to assess the ultrastructure of these membrane bulges and found that they contain layered membrane structures attached to photoreceptor cilia (Figure 1G). To investigate whether these membrane layers retained at the cilium were sufficient to drive CNGβ1 trafficking to the outer segment, we immunostained WT and *Rds*^*-/-*^ retinas for GARP, CNGα1, and ROM1 at P45, when these structures are abundant. Represented by GARP and CNGα1 staining, we observed a dramatic increase in the ciliary localization of the CNG channel in the aged *Rds*^*-/-*^ retinas (Figures 1H and Supplemental Figure 2). Additionally, ROM1 staining is also present in these structures. These findings suggest that in the absence of peripherin-2, the α1- and β1-subunits can form a stable tetramer, but that the channel delivery is enhanced by the accumulation of laminated membranes at the ciliary tip.

### Ciliary delivery of the β1-subunit of the CNG channel is reduced in *Rds*^*-/-*^ mice

While we observed both α1 and β1 subunits in ciliary-derived ectosomes from *Rds*^*-/-*^ mice, it remains unclear why CNG channel levels are specifically reduced. We hypothesized that overexpressed CNGβ1 could drive enhanced ciliary localization of the CNG channel in *Rds*^*-/-*^ rods. MYC-tagged CNGβ1 was overexpressed using a rhodopsin promoter in *Rds*^*-/-*^ rod photoreceptors by *in vivo* electroporation. Overexpression resulted in an abundance of MYC-CNGβ1 expression, but localization was restricted to areas surrounding the nucleus and within internal structures of the inner segment (Figure 2A). This localization pattern suggests that overexpressed CNGβ1 is almost completely trapped in the endomembranes and not in ciliary ectosomes. Importantly, overexpression of other outer segment-resident proteins, such as FLAG-tagged rhodopsin, can traffic to outer segment ectosomes (Figure 2B, (21)). These data suggest the downregulation of CNGβ1 in *Rds*^*-/-*^ rods is a consequence of improper trafficking. We demonstrate this further by co-expressing full-length, FLAG-tagged peripherin-2 with MYC-CNGβ1 in *Rds*^*-/-*^ rods and observe that re-expression of peripherin-2 is sufficient to restore localization of CNGβ1 to the outer segment (Figure 2C). This can also be seen with endogenous CNGβ1 when peripherin-2 is overexpressed and retinal sections are stained for CNGβ1 (Supplementary Figure 3A). CNG channel subunit stoichiometry is important for trafficking, so we confirmed that endogenous CNGα1 could accommodate overexpressed MYC-CNGβ1 by staining with anti-CNGα1 antibodies (Supplementary Figure 3B). We show that this phenomenon is specific to peripherin-2 and not an artifact of overexpression by co-expressing MYC-CNGβ1 and FLAG-rhodopsin in *Rds*^*-/-*^ rods (Figure 2D). FLAG-rhodopsin localizes to ectosomes, but MYC-CNGβ1 remains almost entirely excluded, similar to Figure 2A.

**Figure 2.**
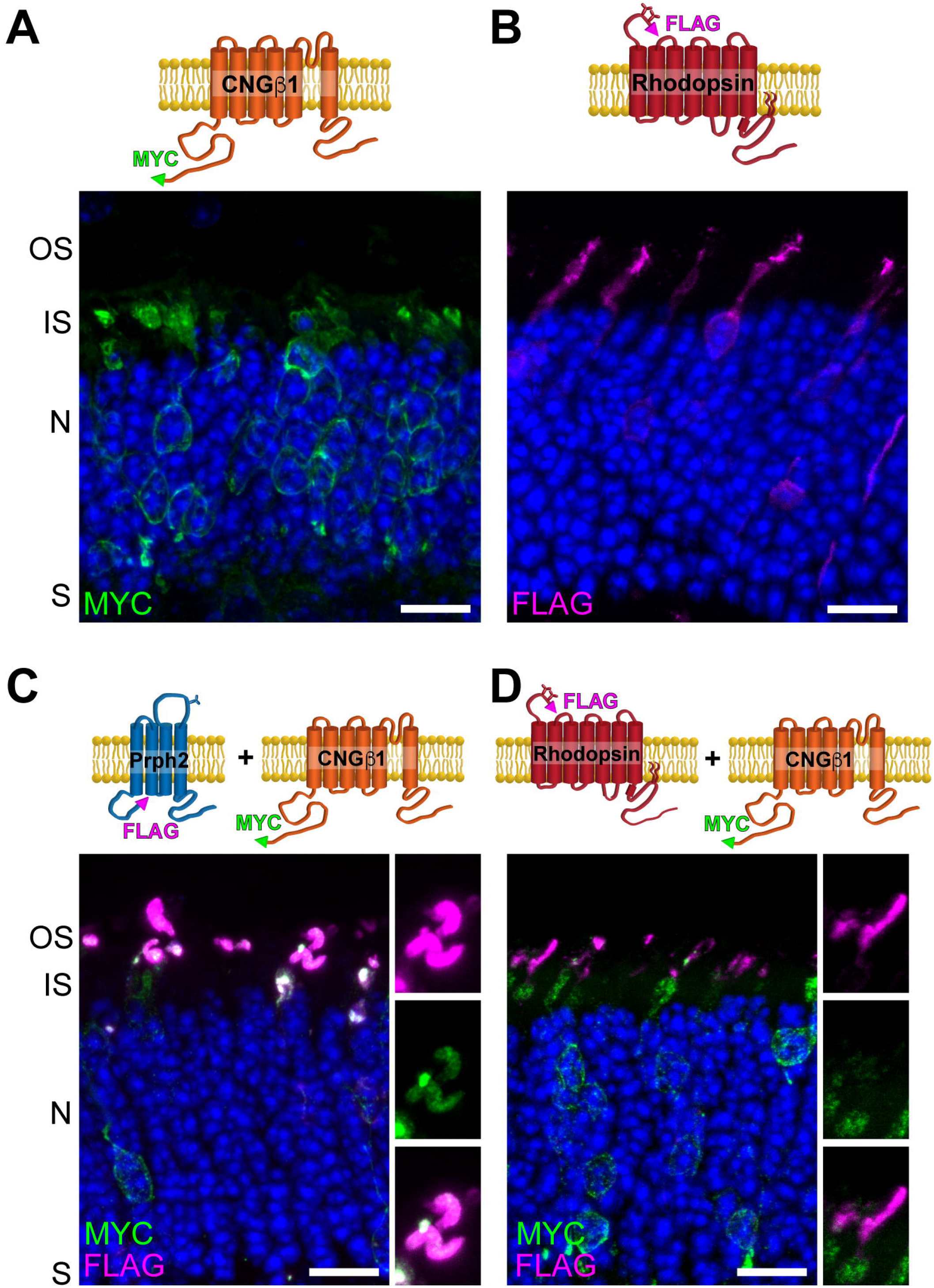
Ciliary delivery of the β1-subunit of the CNG channel is reduced in *Rds*^*-/-*^ mice. *Rds*^*-/-*^ mouse retinal cross-sections expressing **A**. MYC-CNGβ1 (n=3), **B**. FLAG-rhodopsin (n=3), **C**. MYC-CNGβ1 and FLAG-Prph2 (n=5), or **D**. MYC-CNGβ1 and FLAG-Rhodopsin (n=6). Retinal sections were stained for MYC and/or FLAG. Nuclei counterstained in blue. Scale Bar, 10 µm.

### The peripherin-2/ROM1 complex is not required for CNG Channel trafficking to the outer segment

We then wanted to determine how peripherin-2 facilitates trafficking of the CNG channel to the outer segment. One possible scenario could be that peripherin-2 and the CNG channel traffic together to the outer segment. We investigated whether the small proportion of peripherin-2/ROM1 that utilizes the same conventional trafficking route as the CNG channel plays a role in channel delivery. We performed a deglycosylation assay on *Rom1*^*-/-*^ retinal lysates in which all of peripherin-2 traffics unconventionally (20). Wildtype and *Rom1*^*-/-*^ retinal lysates were treated with PNGase F or EndoH to assess the glycosylation state of glycoproteins. PNGase F cleaves all N-linked glycoproteins and is used to monitor the molecular weight shift following deglycosylation. EndoH cannot cleave hybrid oligosaccharide chains that are produced in the Golgi, so proteins that traffic along the conventional pathway, like rhodopsin and CNGα1, are EndoH-resistant. Proteins that rely on the unconventional pathway, like peripherin-2, remain EndoH sensitive (Figure 3A). The CNGβ1 subunit is not glycosylated, so we analyzed the glycosylation state of the CNGα1 subunit because the subunits have been shown to traffic together (13). We confirmed that the fraction of peripherin-2 that traffics conventionally through the Golgi is indeed absent in *Rom1*^*-/-*^ retinas and found that trafficking of the CNG channel *via* the conventional pathway is unaffected (Figure 3A).

**Figure 3.**
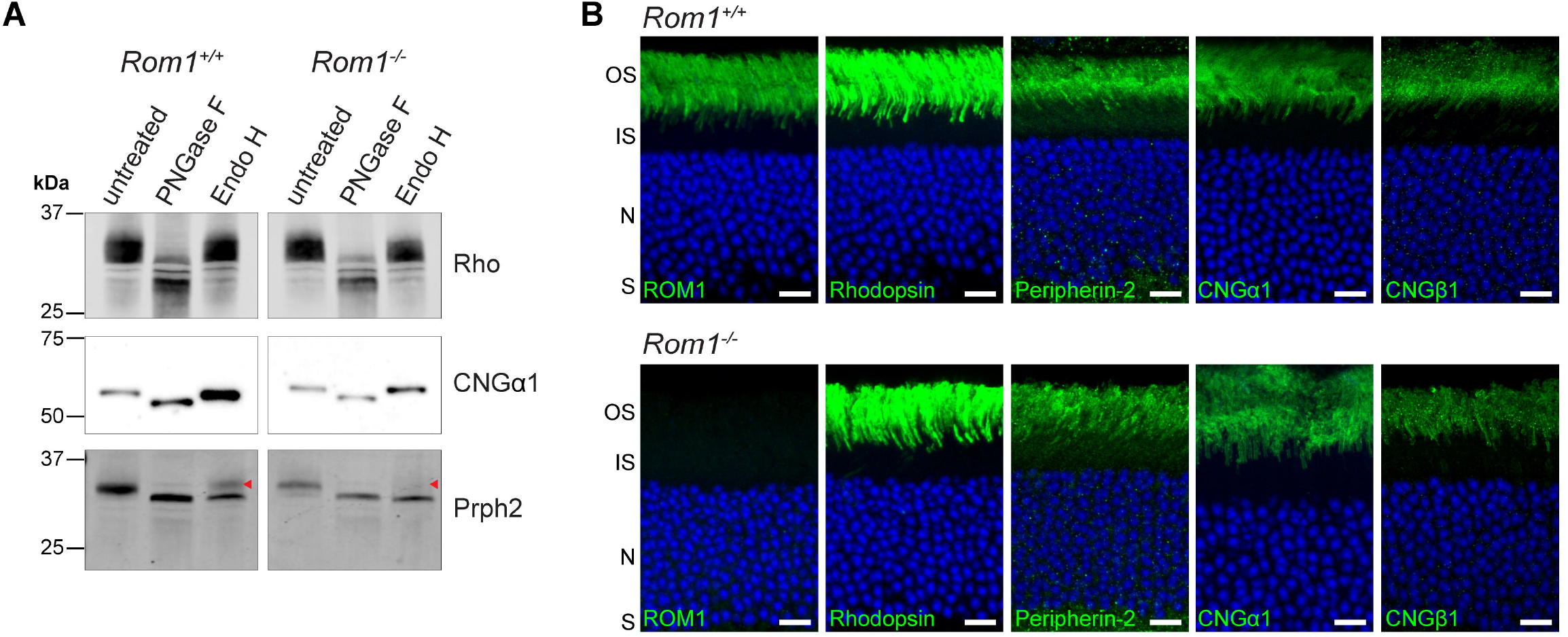
The peripherin-2/ROM1 complex is not required for CNG channel conventional trafficking or localization. **A**. Representative Western blot of *Rom1*^*+/+*^ and *Rom1*^*-/-*^ retinal lysates deglycosylated with either PNGase F or Endo H enzymes and immunoblotted for Rho, CNGα1, or Prph2. Red arrowhead marks EndoH resistant peripherin-2 which is absent in *Rom1*^*-/-*^ **B**. Representative images from *Rom1*^*+/+*^ or *Rom1*^*-/-*^ retinal sections immunostained for ROM1, rhodopsin, peripherin-2, CNGα1, and CNGβ1. Scale Bar, 10 µm. n=3 for all.

*Rom1*^*-/-*^ retinal sections were stained for the channel components, CNGα1 and CNGβ1. We found that both subunits of the CNG channel are properly localized to the plasma membrane of the outer segment in *Rom1*^*-/-*^ retinas (Figure 3B). These data show that CNGβ1 does not require peripherin-2/ROM1 complexes to traffic through the conventional pathway or localize to the outer segment plasma membrane.

### The peripherin tetraspanin membrane domain is sufficient to direct CNGβ1 to the outer segment

Our finding that CNGβ1 transport through the Golgi does not rely on peripherin-2 led us to investigate the role peripherin-2 plays in facilitating outer segment localization of the CNG channel. Specific domains of peripherin-2 were examined in isolation for their ability to restore CNGβ1 localization to the outer segment. We started with two previously characterized fusion proteins that have been shown to localize to the outer segment and build a rudimentary membrane structure (21). The Prph2-Rho_CT_ construct fuses the first 290 amino acids of peripherin-2 – including the four transmembrane domains – to the C-terminal tail of rhodopsin that contains its outer segment targeting motif. Alternatively, the Rho-Prph2_CT_ construct takes the C-terminal, outer-segment targeting region of peripherin-2, and fuses it to the seven transmembrane domains of rhodopsin. We co-expressed these constructs with MYC-CNGβ1 and found that Prph2-Rho_CT_ can rescue MYC-CNGβ1 localization to the outer segment, while Rho-Prph2_CT_ cannot (Figure 4A-C). The percentage of outer segments in *Rds*^*-/-*^ rods with co-localization between MYC-CNGβ1 and FLAG-Prph2 constructs was quantified across animals and displayed as a bar graph in Figure 4F. We confirmed these data by staining for endogenous CNGβ1 in *Rds*^*-/-*^ rods electroporated with each construct (Supplemental Figure 3C,D).

**Figure 4.**
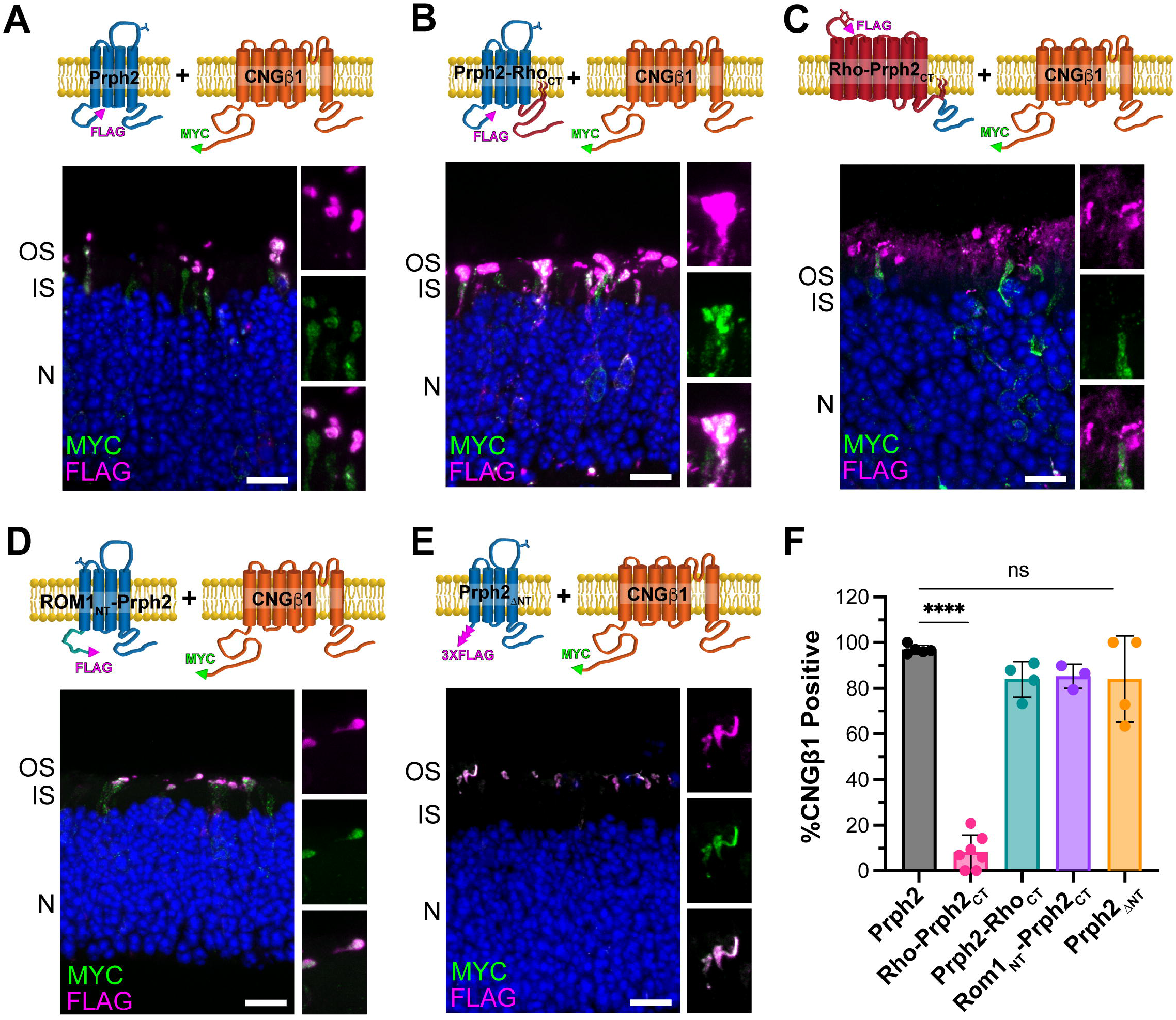
The tetraspanin membrane domain of peripherin-2 is required for CNGβ1 localization to the outer segment. *Rds*^*-/-*^ mouse retinal cross-sections co-expressing MYC-CNGβ1 and FLAG-tagged **A**. Prph2 (n=5), **B**. Prph2-Rho_CT_ (n=4), **C**. Rho-Prph2_CT_ (n=7), **D**. ROM1_NT_-Prph2 (n=3), or **E**. Prph2_ΔNT_ (n=4). Sections are immunostained for MYC and FLAG. Nuclei counterstained in blue. Scale Bar, 10 µm. **F**. Bar graph shows percentage of outer segment structures containing both FLAG (peripherin chimeras) and MYC-CNGβ1. One-way ANOVA was performed with post-hoc t-tests. pValues: ****, p<0.0001; ns, p>0.05.

Rhodopsin was previously shown to facilitate the trafficking of some outer segment membrane proteins (30, 31). To ensure that the trafficking of CNGβ1 is specific to peripherin-2 features, we recreated these chimeras by replacing rhodopsin with a complementary GPCR, the serotonin-6 receptor (Htr6). We fused the C-terminal tail of peripherin-2 to the seven transmembrane domains of Htr6 (Htr6-Prph2_CT_) as well as fused the four transmembrane domains of peripherin-2 to the targeting region of Htr6 (amino acids 392-440, Prph2-Htr6_CT_, (32)). We confirmed that these molecular chimeras can localize to the outer segment in WT retinal sections (Supplemental Figure 4, (21)). Expressing these chimeras in *Rds*^*-/-*^ rods replicates that only the chimera containing the first 290 amino acids of peripherin-2 can rescue CNGβ1 localization to outer segment membranes (Supplemental Figure 5A,B). From these data, we find that the tetraspanin core of peripherin-2 and not its C-terminus restores CNGβ1 localization in *Rds*^*-/-*^ rods.

We then examined the cytosolic N-terminus of peripherin-2 to further narrow down the region of peripherin-2 required for CNGβ1 delivery. Peripherin-2 and ROM1 are structurally homologous, but the amino acid sequences differ within the cytosolic N-terminal region, making ROM1 a good target for a chimeric protein. We started by replacing the N-terminus of peripherin-2 with the N-terminal region of ROM1. When electroporated into *Rds*^*-/-*^ rods we found that MYC-CNGβ1 was localized to the outer segment membranes emanating from the rod cilia (Figure 4D,F). However, we cannot exclude that the N-terminus of ROM1 is redundant to that of peripherin-2, so we followed up by removing peripherin-2’s N-terminus and replacing it with a FLAG-tag (FLAG-Prph2_ΔNT_). After electroporation in WT rods, we found that this construct did not express in the majority of transfected rods and was unable to build rudimentary outer segments in *Rds*^*-/-*^ rods (Supplemental Figure 5C,D). To improve expression, we extended the sequence to a 3XFLAG-tag (3XFLAG-Prph2 _ΔNT_). We found this construct had improved expression in WT rods (Supplemental Figure 4E) and was able to build small, rudimentary outer segments in *Rds*^*-/-*^ rods. When co-expressed with MYC-CNGβ1 in *Rds*^*-/-*^ rods, the 3XFLAG-Prph2_ΔNT_ was sufficient to rescue CNGβ1 localization to these outer segment structures (Figure 4E,F). Overall, these data show that the cytosolic N- and C-termini of peripherin-2 are dispensable for CNGβ1 outer segment localization and reiterate that the tetraspanin region of peripherin-2 is required for CNGβ1 delivery.

### Overexpression of ROM1 constructs in *Rds*^*-/-*^ rods restores CNGβ1 localization to rudimentary outer segments

In a recent study, it was shown that the peripherin-2 homolog, ROM1, can function redundantly to peripherin-2 in outer segment disc formation (33). We were curious if ROM1 is sufficient to facilitate CNGβ1 localization to the outer segment. We started by overexpressing FLAG-ROM1 in *Rds*^*-/-*^ rods and found that FLAG-ROM1 was primarily localized to internal membranes and unable to build ciliary membrane structures (Supplemental Figure 5E), even though it was properly localized to the outer segment in WT rods (Supplemental Figure 3F). Previous studies have improved targeting and expression of ROM1 in *Rds*^*-/-*^ rods by replacing its C-terminus with that of peripherin-2 (20). We co-electroporated MYC-CNGβ1 with a FLAG-tagged ROM1-Prph2_CT_ construct into *Rds*^*-/-*^ rods and found small FLAG-positive structures in the outer segment region that were co-localizing with MYC-CNGβ1 (Figure 5A). We then wanted to determine whether the ROM1 tetraspanin region was sufficient to facilitate CNGβ1 delivery by further replacing ROM1’s cytoplasmic N-terminus with that of peripherin-2 (Prph2_NT/CT_-ROM1). When we co-expressed this construct with MYC-CNGβ1 in *Rds*^*-/-*^ rods we found that Prph2_NT/CT_-ROM1 can rescue MYC-CNGβ1 localization to the outer segment (Figure 5B). Together, this shows that ROM1 is redundant to peripherin-2 in its ability to restore CNGβ1 to the outer segment of *Rds*^*-/-*^ rods and that this function is facilitated through the tetraspanin domain.

**Figure 5.**
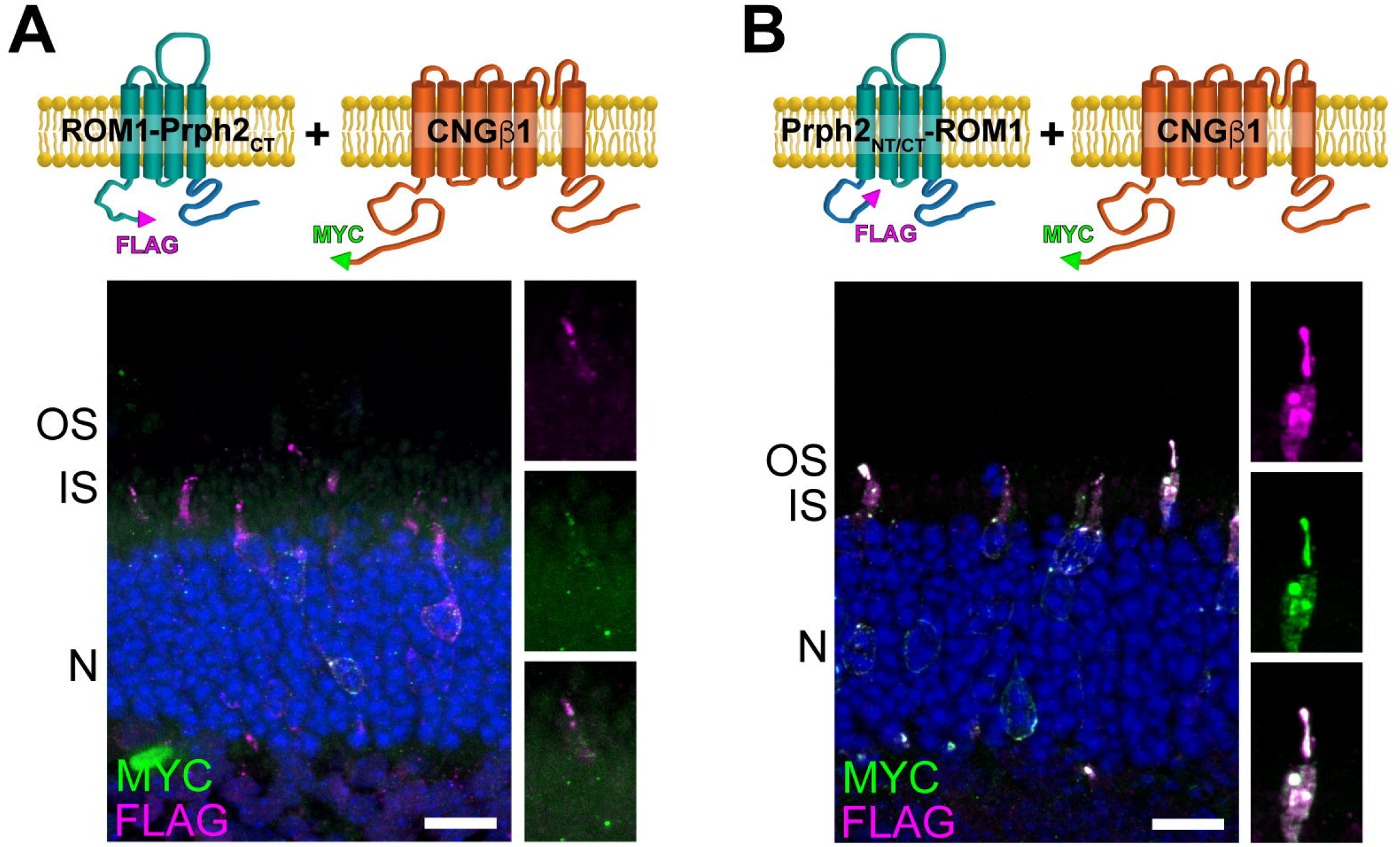
ROM1’s tetraspanin domain restores localization of CNGβ1 to the outer segment. *Rds*^*-/-*^ mouse retinal cross-sections co-expressing MYC-CNGβ1 and FLAG-tagged **A**. ROM1-Prph2_CT_ (n=4) or **B**. Prph2_NT/CT_-ROM1 (n=5). Sections are immunostained for FLAG and MYC. Nuclei counterstained in blue. Scale Bar, 10 µm

## DISCUSSION

The visual response in photoreceptors relies upon proper outer segment organization, which includes the accurate localization of membrane proteins to this organelle. It has been suggested that the structural disc protein, peripherin-2, could guide CNG channel localization to the outer segment. Interactions between these two proteins have been identified, and it was reported that CNGβ1’s localization was compromised in the absence of peripherin-2. The mechanism behind the CNG channel’s dependence on peripherin-2 to make it to the outer segment was perplexing, as peripherin-2 and CNGβ1 mostly utilize separate trafficking pathways to reach the outer segment and reside in two distinct membrane domains within the outer segment. We set out to investigate how peripherin-2 may be involved in the localization of the CNG channel to the photoreceptor outer segment.

We found in *Rds*^*-/-*^ retinas that mRNA levels of several outer segment genes are reduced. We posit that the loss of the outer segment compartment drives general downregulation of outer segment genes. We noted that the amount of CNG channel in the outer segment region in *Rds*^*-/-*^ retina sections from young mice was severely reduced, which is consistent with previous reports (22, 23). When we examined protein levels, we found that peripherin-2’s binding partner, ROM1, and the CNG channel subunits were severely downregulated when compared to the disc protein ABCA4. In fact, only when we examined a concentrated preparation of ciliary-derived ectosomes from these mice did we observe that low levels of the CNG channel are actually present. Our finding that there is a decrease in the CNG channel at the protein level suggests that not all outer segment proteins behave similarly when the outer segment is not formed. Supporting this notion, previous reports have shown the reverse effect: PCHD21 protein is upregulated in *Rds*^*-/-*^ retinas (34). The selective downregulation of the CNG channel in the absence of peripherin-2 is curious, as other models of outer segment dysmorphia (e.g., *Rho*^*-/-*^, *PCDH21*^*-/-*^, or *WASF3*^*-/-*^) localize the CNG channel properly (30, 35, 36). Recent studies have investigated mechanisms driving G protein-coupled receptor sorting into extracellular vesicles released from the ciliary membrane and inner segment plasma membrane (42-44). It is possible that these or similar mechanisms play a role in excluding the CNG channel from ciliary ectosomes, which could result in its selective downregulation.

Our study went on to examine factors that contribute to CNG channel ciliary localization. One hypothesis was that peripherin-2 facilitates CNG channel trafficking to the outer segment – possibly through the small fraction of peripherin-2 that traffics through the Golgi in a complex with ROM1. We examined *ROM1*^*-/-*^ retinas, in which no peripherin-2 goes through the Golgi, and found CNG channel trafficking to the outer segment was unaffected. These data show that peripherin-2 and the CNG channel traffic separately, and are corroborated by previous work from our lab that found that CNGβ1 outer segment targeting occurs in the absence of peripherin-2 binding (13). While our data do not rule out that CNGβ1 and peripherin-2 could interact before trafficking pathways diverge, we find this interaction does not play a role in CNG channel delivery to the cilia. We went on to determine the minimal region of peripherin-2 that can restore CNGβ1 delivery to the outer segment. We originally hypothesized that the C-terminus of peripherin-2 would be critical for CNGβ1 delivery because this region contains both the outer segment targeting sequence for peripherin-2 and a signal to retain ectosomes at the ciliary tip (21, 37, 38). However, we found that both the cytoplasmic N- and C-terminal regions were dispensable, and rather the membrane-spanning domain is necessary and sufficient for CNGβ1 delivery.

The major role of the tetraspanin domain of peripherin-2 is to create curvature in the membrane through its assembly into homotetramers or heterotetramers with its homolog, ROM1. These tetramers are fundamental units that create the hairpin structure at the edges of outer segment discs known as the rim (39, 40). The tetraspanin proteins peripherin-2 and ROM1 are also required in proper ratios for successful disc enclosure (33, 41). The essential role that these tetraspanins play in forming disc membranes suggests the delivery of the CNG channel may be driven by the delineation of a lamellar membrane within the outer segment. In the P45 *Rds*^*-/-*^ retinas, we found that ciliary protrusions also contain ROM1, which has been shown to have redundant functions with peripherin-2 (33). We hypothesize that lamellar protrusions in *Rds*^*-/-*^ at P45 are built by ROM1; however, we were unable to increase their prevalence through overexpression of FLAG-tagged full-length ROM1. We were able to circumvent this by appending the C-terminus of peripherin-2 to the tetraspanin domain of ROM1, which we found was sufficient to restore CNGβ1.

In the photoreceptor outer segment, discs are formed as an evagination of the ciliary membrane through a lamellipodia-like mechanism that relies on branched actin filaments. It has been shown that inhibiting actin polymerization prevents new disc formation (26) and also prevents CNG channel entry into the plasma membrane (45). These data suggest that these two events are linked. Interestingly, when actin polymerization is inhibited, the newly synthesized CNG channel is held at the distal end of the connecting cilium, primed for entry (45). This is also the site where tetraspanin proteins are found during new disc synthesis as they are restricted to the axonemal side of nascent discs (45-48). Together, these data suggest that the partitioning of disc membranes from the plasma membrane by tetraspanin proteins is a key step for localizing the CNG channel and could play a role in segregating other proteins into the plasma membrane. Our results support this conclusion, and additional evidence comes from analysis of rhodopsin knockout outer segments, which have internal disc-like structures and localize the CNG channel properly (30). Therefore, peripherin-2’s ability to delineate a disc is sufficient to drive CNG channel localization even in the absence of a fully elaborated outer segment.

## METHODS

### Animals

*Rds*^*-/-*^ described by (28) and confirmed KO by (49). WT *CD1* mice used in electroporation experiments were obtained from Charles River; WT *C57BL/6J* and *Rds*^*-/-*^ from The Jackson Laboratory. *Rds*^*-/-*^ mice maintained in our colony were backcrossed to *C57BL/6J* to produce *Rds*^*+/-*^ and *Rds*^*+/+*^ littermates, included in qRT-PCR, immunofluorescence, and Western blot experiments as heterozygous and wildtype mice, respectively. All animals were analyzed at P21 unless otherwise noted. Mice were handled following protocols approved by the Institutional Animal Care and Use Committee at the University of Michigan (registry number A3114-01). As there are no known gender-specific differences in vertebrate photoreceptor development and/or function, male and female mice were randomly assigned to experimental groups. All mice were housed in a 12/12-hr light/dark cycle with free access to food and water.

### DNA constructs

DNA constructs were generated using overhang extension PCR methods. Generation of Prph2-Rho_CT_, Rho-Per_CT_, and Htr6-Per_CT_ was previously described (21). Primers used to generate remaining molecular fusion proteins are detailed in Supplementary Table 2. All forward and reverse primers were designed to introduce AgeI and NotI restriction digest sites, respectively. For rod-specific expression, cDNA sequences were cloned between 5’ *AgeI* and 3’ *NotI* cloning sites into the pRho vector to drive expression by a 2.2 kb bovine rhodopsin promoter (Addgene plasmid #11156; http://n2t.net/addgene:11156).

### In vivo *electroporation*

DNA constructs were electroporated into neonatal mouse retinas using the protocol described in (50) with modifications detailed in (21). Briefly, neonatal (P0-P2) mice were anesthetized on ice, followed by puncture of the eyelid and cornea with a 30-gauge needle at the periphery of the eye. A blunt-end, 32-gauge Hamilton syringe was used to inject 0.3-0.5 µL of concentrated plasmid DNA subretinally (∼2 µg/µL of construct of interest mixed with ∼1 µg/µL of mCherry transfection marker). The positive side of a tweezer-type electrode was placed over the injected eye (BTX, Holliston, MA) and five 50 ms pulses were applied at 100-110 V using an ECM830 square pulse generator (BTX). After neonates recover, they are returned to their mother to develop until collection at P20-P22.

### Immunofluorescence

Mouse eyes were collected and drop-fixed with 4% paraformaldehyde in 1X PBS (Fisher Scientific, AAJ75889K8) for 1-2 hrs at room temperature. Eyes were washed and stored in 1X PBS before they were dissected into posterior eyecups. Eyecups were embedded in 4% agarose (Fisher Scientific, BP160-500) and cut into 100 µm sections through the central retina using a Leica VT1200s vibratome. Floating sections were blocked in 5% donkey serum and 0.5% Triton X-100 in 1X PBS for 1 hr at room temperature before incubation with primary antibody overnight at 4ºC. The following day, sections were washed with 1X PBS and then incubated with fluorophore-conjugated secondary antibodies and 10 µg/ml DAPI (Sigma-Aldrich, 102362760) for 1-2 hrs at room temperature. Sections were then rinsed three times in 1X PBS and mounted with Prolong Glass (Thermo Fisher Scientific, P36980) under High Precision #1.5 coverslips (Electron Microscopy Sciences, 72204-10). Images were acquired using a Zeiss Observer 7 inverted microscope equipped with a 63X oil-immersion objective (1.40 NA) and LSM 800 confocal scanhead controlled by Zen 5.0 software (Zeiss).

### Enriched Ectosome Preparation

Ectosomes were purified from *Rds*^*-/-*^ mice using a protocol adapted from (26). In brief, at least ten *Rds*^*-/-*^ retinas were detached in ice cold Ringer’s solution containing 130 mM NaCl, 3.6 mM KCl, 2.4 mM MgCl2, 1.2 mM CaCl2, and 10 mM Hepes (pH 7.4) solution containing Complete Protease Inhibitor (CPI; Millipore Sigma, 11836170001). Detached retinas were transferred to a 1.5mL low-retention microcentrifuge tube and agitated at 4 ºC for 10 minutes. Large debris was removed by centrifugation at 300 g for 5 min at 4 ºC. The supernatant was further cleared by centrifugation at 2000 g for 5 min at 4 ºC. The ectosome fraction was then pelleted by ultracentrifugation at 150,000 g for 20 min at 4 ºC. The membrane pellet was resuspended in 20 µL of 2% SDS in PBS+CPI. Multiple volumes (10, 5, 2.5, and 1.25 µL) of the sample was then loaded on an SDS-PAGE gel followed by Western blot.

### Western Blot and Deglycosylation

Whole retinas were sonicated in 200 µL of 1X PBS with CPI and centrifuged at 60,000 rpm for 20 minutes at 4 ºC. The supernatant was collected as the soluble fraction, and the pellet was resuspended in 250 µL of 2% SDS in PBS with CPI and collected as the membrane fraction. Membrane fractions were subsequently spun at 14,000 rpm to pellet any remaining insoluble material. A DC Assay was performed on the cleared membrane fractions and compared to a BSA standard curve to determine protein concentration. Samples were diluted to a concentration 0.8-1 µg/µL in Laemelli sample buffer with 5% β-mercaptoethanol and loaded on an AnyKd Mini-PROTEAN TGX Precast Protein Gels (Bio-Rad, 4569033) in increasing µL amounts to achieve the desired µg amount. SDS-PAGE gels were run at 50 V for 30 min before increasing to 120 V until the dye front ran to the bottom (∼1 hr).

The SDS-PAGE gel was transferred to Immun-Blot Low Fluorescence PVDF Membrane (Bio-Rad, 1620264) at 90 mV for 90 minutes. Membranes were blocked for ∼1 hr with Intercept Blocking Buffer (LI-COR Biosciences, 927-70003) at room temperature before incubation with primary antibodies diluted in 1:1 ratio of Blocking Buffer and PBS-T overnight at 4ºC. The next day, membranes were washed with 1X PBS-T and incubated with IRDye secondary antibodies diluted in 1:1 ratio of Blocking Buffer and PBS-T with 0.2% SDS for 1-2 hrs at room temperature. Bands were visualized and quantified using the Odyssey CLx infrared imaging system (LiCor Bioscience). Images of the uncropped Western blots can be found in Supplementary Figure 6. For CNGα1 immunoblots, membranes were blocked for ∼1 hr with 5% non-fat dry milk, 5% donkey serum, and 1% TritonX-100 in PBS at room temperature before incubation with primary antibodies in blocking solution overnight at 4ºC. The next day, membranes were washed with 1X PBS-T and incubated with an HRP-conjugated secondary antibody diluted in blocking solution for 1-2 hrs at room temperature. Bands were visualized and quantified using the Azure imaging system at multiple exposure times ranging from 30 s to 20 min.

To perform a deglycosylation assay, whole retinal lysates were prepared by sonicating one retina in 250 μL of 2% SDS in 1X PBS with CPI. Lysates were cleared by centrifugation at 14,000 rpm and 23ºC. Protein concentration was then determined by DC Assay and three 20 µL reactions containing 10 µg of retinal lysate were prepared: control, PNGase F, and EndoH (NEB, P0708L and P0702L). Control and PNGase F reactions contained 2 µL GlycoBuffer 2, 2 µL of 10% NP-40, and 2 µL of PNGase F or water, respectively. The EndoH reaction contained 2 µL GlycoBuffer 3 and 2 µL EndoH. Reactions were incubated for 1 hr at 37ºC before performing the Western Blot procedure as described above.

### Antibodies

The following commercial primary antibodies were used: mouse monoclonal M2 anti-FLAG (Sigma-Aldrich, F1804); rabbit polyclonal anti-FLAG (Sigma-Aldrich, F7425); goat polyclonal anti-FLAG (Abcam, ab1257); rabbit monoclonal 71D10 anti-MYC (Cell Signaling, 2278S); mouse monoclonal 9B11 anti-MYC (Cell Signaling, 2276S); mouse monoclonal 1D4 anti-rhodopsin (Abcam, ab5417); goat polyclonal M-18 anti-ABCA4 (Santa Cruz, SC-21460); mouse monoclonal PMc 1D1 anti-CNGα1 (Abcam, ab255766); goat polyclonal anti-ABCA4 (Everest Biotech, EB08615). The following antibodies were generously provided by: Vadim Arshavsky, Duke University (sheep polyclonal anti-ROM-1); Robert Molday, University of British Columbia (mouse monoclonal 1D1 PMC, anti-CNGα1; mouse monoclonal 4B1, anti-GARP); Steven Pittler, University of Alabama at Birmingham (rabbit polyclonal anti-CNGβ1).

The following commercial secondary antibodies were used: Donkey anti-mouse AF488 (A21202, Fisher Scientific); Donkey anti-rabbit AF488 (A21206, Fisher Scientific); Donkey anti-sheep (A11015, Fisher Scientific); Donkey anti-sheep AF568 (A21099, Fisher Scientific); Donkey anti-mouse AF647 (A31571, Fisher Scientific); Donkey anti-Sheep AF647 (A21448, Fisher Scientific); Donkey anti-mouse HRP (Jackson ImmunoResearch 715-035-150).

### Scanning electron microscopy

Mouse retinas were dissected in ice cold Ringer’s solution adjusted to 314 mOsm. The retinas were fixed with 2% paraformaldehyde, 2% glutaraldehyde, and 0.05% calcium chloride in 50 mM MOPS (pH 7.4) for 1 hour with gentle agitation. Retinas were then washed with PBS and post-fixed with 1% Osmium Tetroxide in PBS (Electron Microscopy Sciences – EMS) for 1 hour. Next, the retinas were washed with PBS, subsequently dehydrated through graded ethanol to 100% and finally dried via the critical point method utilizing a Tousimis Samdri PVT-3D. Dried retinas were then mounted onto 12mm SEM stubs with carbon conductive paint (EMS) and immediately sputter coated with ∼15 nm of Au-Pd alloy utilizing a Denton Desk-V coater. Lastly, retinas were analyzed with a JEOL JSM-IT100 Scanning Electron Microscope operated at 10kV, and digital micrographs collected with an Everhart-Thornley secondary electron detector.

### Transmission electron microscopy

Fixation and processing of mouse eyes for thin plastic sections was performed as described previously (47). Mice were deeply anesthetized and transcardially perfused with 2% paraformaldehyde, 2% glutaraldehyde, and 0.05% calcium chloride in 50 mM MOPS (pH 7.4) resulting in exsanguination. The eyes were enucleated and fixed for an additional 2 h in the same buffer at 22 °C. The fixed eyes were washed in PBS, eyecups were dissected and embedded in PBS containing 2.5% agarose (KU VF-AGT-VM; Precisionary), and cut into 200 µm thick slices on a vibratome (VT1200S; Leica). The vibratome sections were stained with 1% tannic acid (Electron Microscopy Sciences) and 1% uranyl acetate (Electron Microscopy Sciences), gradually dehydrated with ethanol and embedded in Spurr’s resin (Electron Microscopy Sciences). 70 nm sections were cut, placed on copper grids, and counterstained with 2% uranyl acetate and 3.5% lead citrate (19314; Ted Pella). The samples were imaged on a JEM-1400 electron microscope (JEOL) at 60 kV with a digital camera (Orius SC200D; Gatan).

## Acknowledgments

The ROM1^-/-^ and littermate control eyes were generously provided by Dr. Andrew Goldberg (Oakland University). The authors would also like to thank Ben Zink at SUNY Upstate Medical University for technical expertise in transmission electron microscopy and scanning electron microscopy.

## REFERENCES

1. R. W. Young, D. Bok, Participation of the retinal pigment epithelium in the rod outer segment renewal process. J. Cell Biol. 42, 392–403 (1969).

2. R. W. Young, The renewal of photoreceptor cell outer segments. J. Cell Biol. 33, 61–72 (1967).

3. D. S. Williams, Transport to the photoreceptor outer segment by myosin VIIa and kinesin II. Vision Res. 42, 455–462 (2002).

4. J. N. Pearring, R. Y. Salinas, S. A. Baker, V. Y. Arshavsky, Protein sorting, targeting and trafficking in photoreceptor cells. Prog. Retin. Eye Res. 36, 24–51 (2013).

5. N. P. Skiba et al., Proteomic identification of unique photoreceptor disc components reveals the presence of PRCD, a protein linked to retinal degeneration. J Proteome Res 12, 3010–3018 (2013).

6. N. P. Skiba et al., TMEM67, TMEM237, and Embigin in Complex With Monocarboxylate Transporter MCT1 Are Unique Components of the Photoreceptor Outer Segment Plasma Membrane. Mol Cell Proteomics 20, 100088 (2021).

7. R. S. Molday, L. L. Molday, Differences in the protein composition of bovine retinal rod outer segment disk and plasma membranes isolated by a ricin-gold-dextran density perturbation method. J. Cell Biol. 105, 2589–2601 (1987).

8. A. Poetsch, L. L. Molday, R. S. Molday, The cGMP-gated channel and related glutamic acid-rich proteins interact with peripherin-2 at the rim region of rod photoreceptor disc membranes. J. Biol. Chem. 276, 48009–48016 (2001).

9. D. C. A. Barret, U. B. Kaupp, J. Marino, The structure of cyclic nucleotide-gated channels in rod and cone photoreceptors. Trends Neurosci 45, 763–776 (2022).

10. J. Xue, Y. Han, W. Zeng, Y. Jiang, Structural mechanisms of assembly, permeation, gating, and pharmacology of native human rod CNG channel. Neuron 110, 86–95.e85 (2022).

11. U. B. Kaupp et al., Primary structure and functional expression from complementary DNA of the rod photoreceptor cyclic GMP-gated channel. Nature 342, 762–766 (1989).

12. T. Y. Chen et al., A new subunit of the cyclic nucleotide-gated cation channel in retinal rods. Nature 362, 764–767 (1993).

13. J. N. Pearring et al., The GARP Domain of the Rod CNG Channel’s β1-Subunit Contains Distinct Sites for Outer Segment Targeting and Connecting to the Photoreceptor Disk Rim. J Neurosci 41, 3094–3104 (2021).

14. Y. Zhang et al., Knockout of GARPs and the β-subunit of the rod cGMP-gated channel disrupts disk morphogenesis and rod outer segment structural integrity. J Cell Sci 122, 1192–1200 (2009).

15. S. Huttl et al., Impaired channel targeting and retinal degeneration in mice lacking the cyclic nucleotide-gated channel subunit CNGB1. J. Neurosci. 25, 130–138 (2005).

16. G. Tian et al., An unconventional secretory pathway mediates the cilia targeting of peripherin/rds. J. Neurosci. 34, 992–1006 (2014).

17. D. Deretic, D. S. Papermaster, Polarized sorting of rhodopsin on post-Golgi membranes in frog retinal photoreceptor cells. J. Cell Biol. 113, 1281–1293 (1991).

18. G. J. Connell, R. S. Molday, Molecular cloning, primary structure, and orientation of the vertebrate photoreceptor cell protein peripherin in the rod outer segment disk membrane. Biochemistry 29, 4691–4698 (1990).

19. M. Illing, L. L. Molday, R. S. Molday, The 220-kDa rim protein of retinal rod outer segments is a member of the ABC transporter superfamily. J Biol Chem 272, 10303–10310 (1997).

20. S. M. Conley et al., Prph2 initiates outer segment morphogenesis but maturation requires Prph2/Rom1 oligomerization. Hum. Mol. Genet. 28, 459–475 (2019).

21. R. Y. Salinas et al., Photoreceptor discs form through peripherin-dependent suppression of ciliary ectosome release. J. Cell Biol. 216, 1489–1499 (2017).

22. D. Chakraborty, S. M. Conley, M. L. DeRamus, S. J. Pittler, M. I. Naash, Varying the GARP2-to-RDS Ratio Leads to Defects in Rim Formation and Rod and Cone Function. Invest Ophthalmol Vis Sci 56, 8187–8198 (2015).

23. D. Chakraborty, S. M. Conley, S. J. Pittler, M. I. Naash, Role of RDS and Rhodopsin in Cngb1-Related Retinal Degeneration. Invest Ophthalmol Vis Sci 57, 787–797 (2016).

24. L. M. Ritter et al., In situ visualization of protein interactions in sensory neurons: glutamic acid-rich proteins (GARPs) play differential roles for photoreceptor outer segment scaffolding. J. Neurosci. 31, 11231–11243 (2011).

25. M. D. Ardell, D. L. Bedsole, R. V. Schoborg, S. J. Pittler, Genomic organization of the human rod photoreceptor cGMP-gated cation channel beta-subunit gene. Gene 245, 311–318 (2000).

26. W. J. Spencer et al., Photoreceptor disc membranes are formed through an Arp2/3-dependent lamellipodium-like mechanism. Proc Natl Acad Sci U S A 116, 27043–27052 (2019).

27. A. I. Cohen, Some cytological and initial biochemical observations on photoreceptors in retinas of rds mice. Invest. Ophthalmol. Vis. Sci. 24, 832–843 (1983).

28. S. Sanyal, H. G. Jansen, Absence of receptor outer segments in the retina of rds mutant mice. Neurosci. Lett. 21, 23–26 (1981).

29. H. G. Jansen, S. Sanyal, W. J. De Grip, J. J. Schalken, Development and degeneration of retina in rds mutant mice: ultraimmunohistochemical localization of opsin. Exp Eye Res 44, 347–361 (1987).

30. J. N. Pearring, W. J. Spencer, E. C. Lieu, V. Y. Arshavsky, Guanylate cyclase 1 relies on rhodopsin for intracellular stability and ciliary trafficking. eLife 4 :e12058 (2015).

31. W. J. Spencer et al., Progressive Rod-Cone Degeneration (PRCD) Protein Requires N-Terminal S-Acylation and Rhodopsin Binding for Photoreceptor Outer Segment Localization and Maintaining Intracellular Stability. Biochemistry 55, 5028–5037 (2016).

32. P. Barbeito et al., HTR6 and SSTR3 ciliary targeting relies on both IC3 loops and C-terminal tails. Life Sci Alliance 4 (2021).

33. T. R. Lewis et al., ROM1 is redundant to PRPH2 as a molecular building block of photoreceptor disc rims. Elife 12 (2023).

34. A. Rattner, J. Chen, J. Nathans, Proteolytic shedding of the extracellular domain of photoreceptor cadherin. Implications for outer segment assembly. J Biol Chem 279, 42202–42210 (2004).

35. A. Rattner et al., A photoreceptor-specific cadherin is essential for the structural integrity of the outer segment and for photoreceptor survival. Neuron 32, 775–786 (2001).

36. W. J. Spencer et al., The WAVE complex drives the morphogenesis of the photoreceptor outer segment cilium. Proc Natl Acad Sci U S A 120, e2215011120 (2023).

37. R. Y. Salinas, S. A. Baker, S. M. Gospe, V. Y. Arshavsky, A single valine residue plays an essential role in peripherin/rds targeting to photoreceptor outer segments. PLoS One 8, e54292 (2013).

38. B. M. Tam, O. L. Moritz, D. S. Papermaster, The C terminus of peripherin/rds participates in rod outer segment targeting and alignment of disk incisures. Mol. Biol. Cell 15, 2027–2037 (2004).

39. M. Poge et al., Determinants shaping the nanoscale architecture of the mouse rod outer segment. Elife 10 (2021).

40. M. L. Milstein et al., Multistep peripherin-2/rds self-assembly drives membrane curvature for outer segment disk architecture and photoreceptor viability. Proc. Natl. Acad. Sci. U. S. A. 117, 4400–4410 (2020).

41. T. R. Lewis et al., Photoreceptor disc incisures form as an adaptive mechanism ensuring the completion of disc enclosure. Elife 12 (2023).

42. A. R. Nager et al., An Actin Network Dispatches Ciliary GPCRs into Extracellular Vesicles to Modulate Signaling. Cell 168, 252–263.e214 (2017).

43. T. R. Lewis et al., Contribution of intraflagellar transport to compartmentalization and maintenance of the photoreceptor cell. Proc Natl Acad Sci U S A 121, e2408551121 (2024).

44. P. Ropelewski, Y. Imanishi, RPE Cells Engulf Microvesicles Secreted by Degenerating Rod Photoreceptors. eNeuro 7 (2020).

45. I. Nemet, G. Tian, Y. Imanishi, Submembrane assembly and renewal of rod photoreceptor cGMP-gated channel: insight into the actin-dependent process of outer segment morphogenesis. J. Neurosci. 34, 8164–8174 (2014).

46. N. V. Klementieva, T. R. Lewis, O. Alekseev, V. Y. Arshavsky, Peripherin-2 and ROM1 Incorporate Directly Into the Rims of Enclosing Photoreceptor Discs Without Accumulating in the Nascent Disc Lamellae. Invest Ophthalmol Vis Sci 66, 33 (2025).

47. J.-D. Ding, R. Y. Salinas, V. Y. Arshavsky, Discs of mammalian rod photoreceptors form through the membrane evagination mechanism. J Cell Biol 211, 495–502 (2015).

48. T. Burgoyne et al., Rod disc renewal occurs by evagination of the ciliary plasma membrane that makes cadherin-based contacts with the inner segment. Proc Natl Acad Sci U S A 112, 15922–15927 (2015).

49. J. Ma et al., Retinal degeneration slow (rds) in mouse results from simple insertion of a t haplotype-specific element into protein-coding exon II. Genomics 28, 212–219 (1995).

50. T. Matsuda, C. L. Cepko, Controlled expression of transgenes introduced by in vivo electroporation. Proc Natl Acad Sci USA 104, 1027–1032 (2007).

