## Supplemental Figures for "The tetraspanin disc proteins, peripherin-2 and ROM1, facilitate CNG channel localization to the rod outer segment"

**Supplementary Figures**

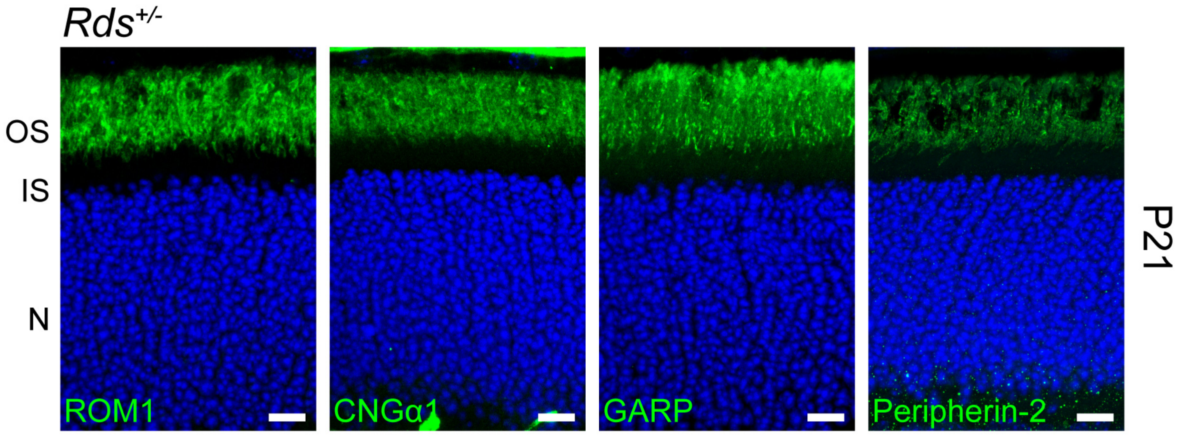


**Supplementary Figure 1.** Retinal sections from WT and *Rds*$^{-/-}$ mice immunostained for ROM1, CNGα1, GARP proteins, and peripherin-2. Nuclei counterstained in blue. n=3 for all. Scale Bar, 10 µm.

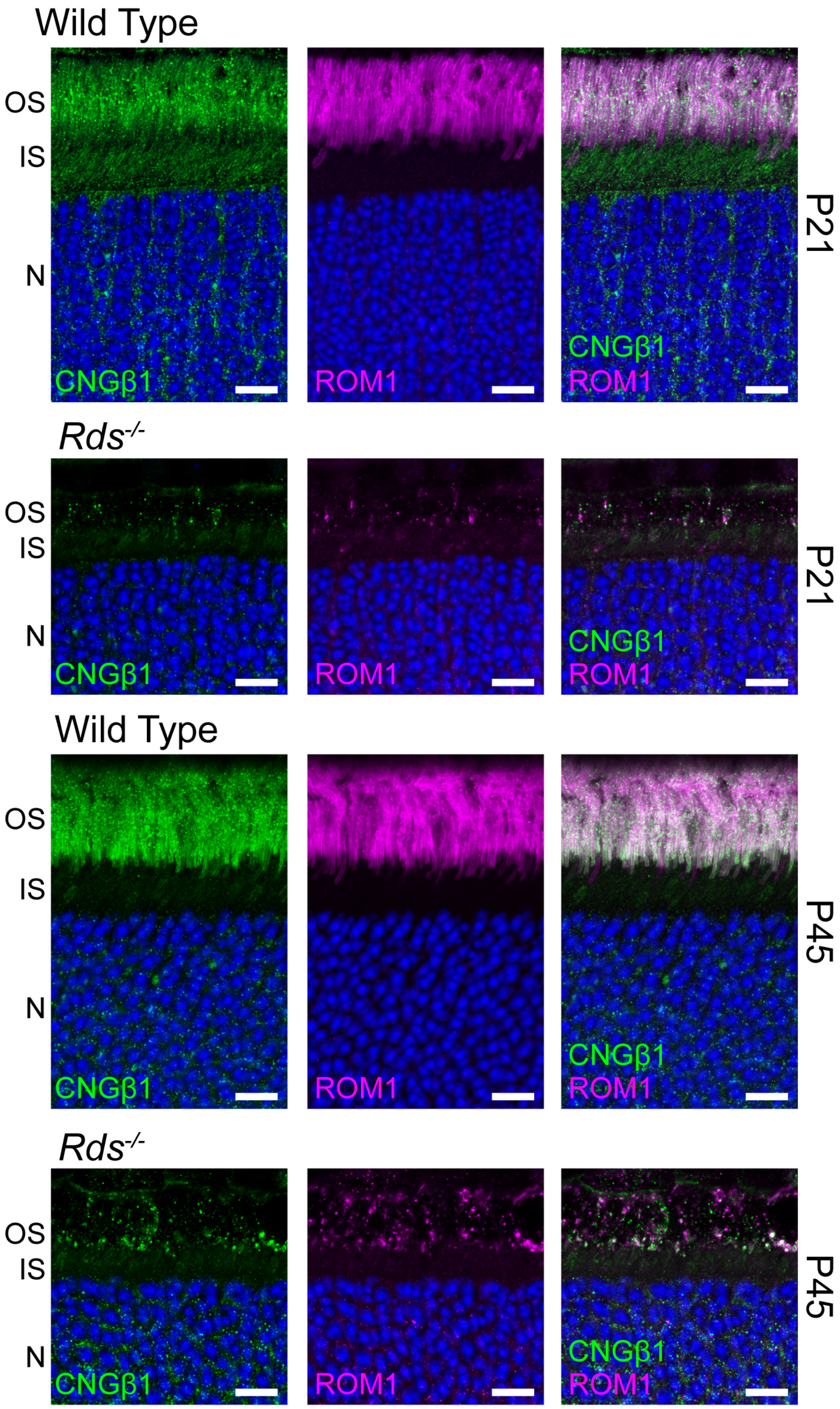


**Supplementary Figure 2.** Retinal sections from WT and *Rds*$^{-/-}$ mice at P21 and P45 immunostained for CNGβ1 and ROM1. Nuclei counterstained in blue. Scale Bar, 10 μm.

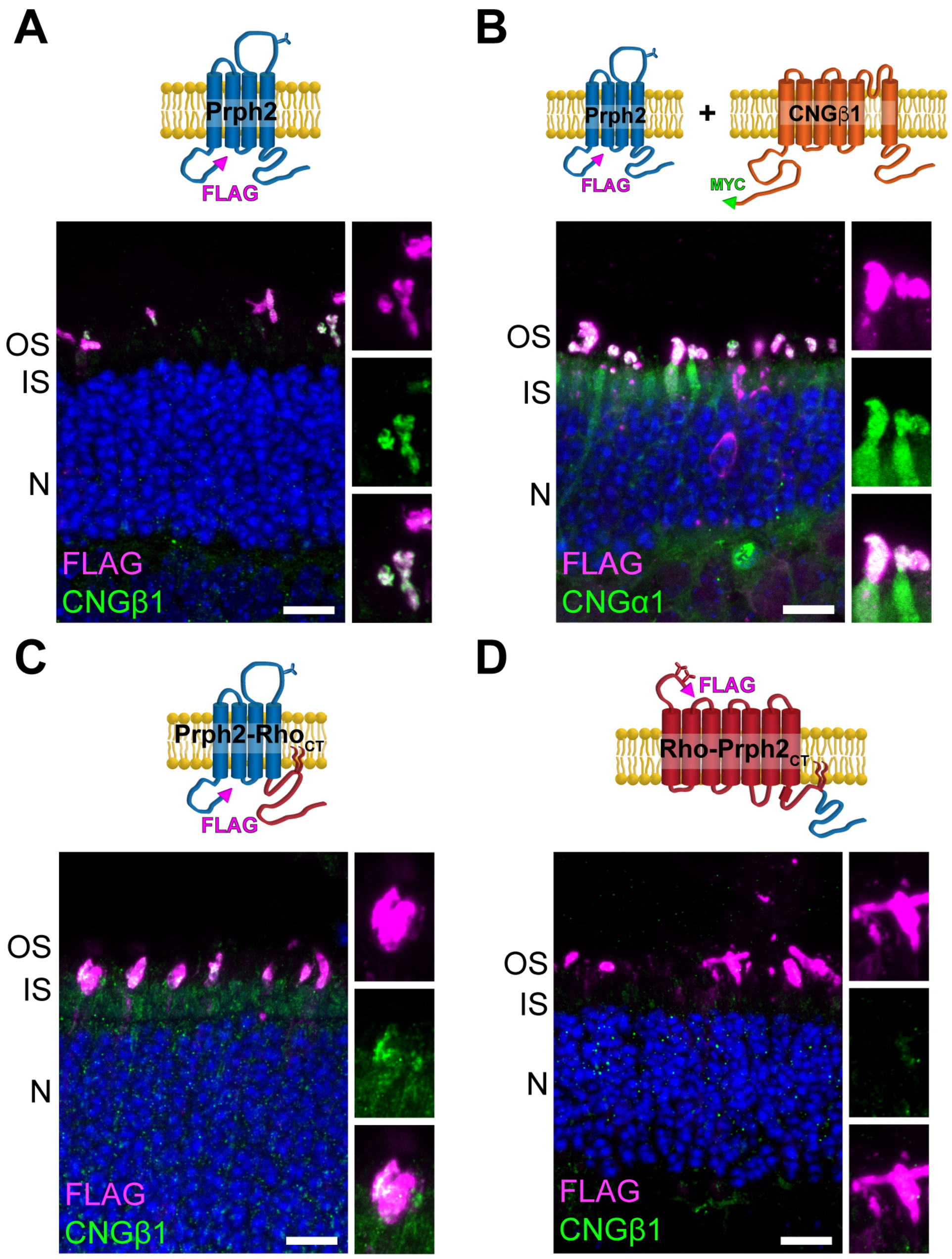


**Supplementary Figure 3.** *Rds*$^{-/-}$ mouse retinal cross-sections expressing **A.** FLAG-Prph2 (n=5), **B.** FLAG-Prph2 and MYC-CNGβ1 **C.** Prph2-Rho$_{CT}$, **D.** Rho-Prph2$_{CT}$ (n=7). Retinal sections immunostained for FLAG, CNGβ1, or CNGα1. Nuclei counterstained in blue. Scale Bar, 10 μm.

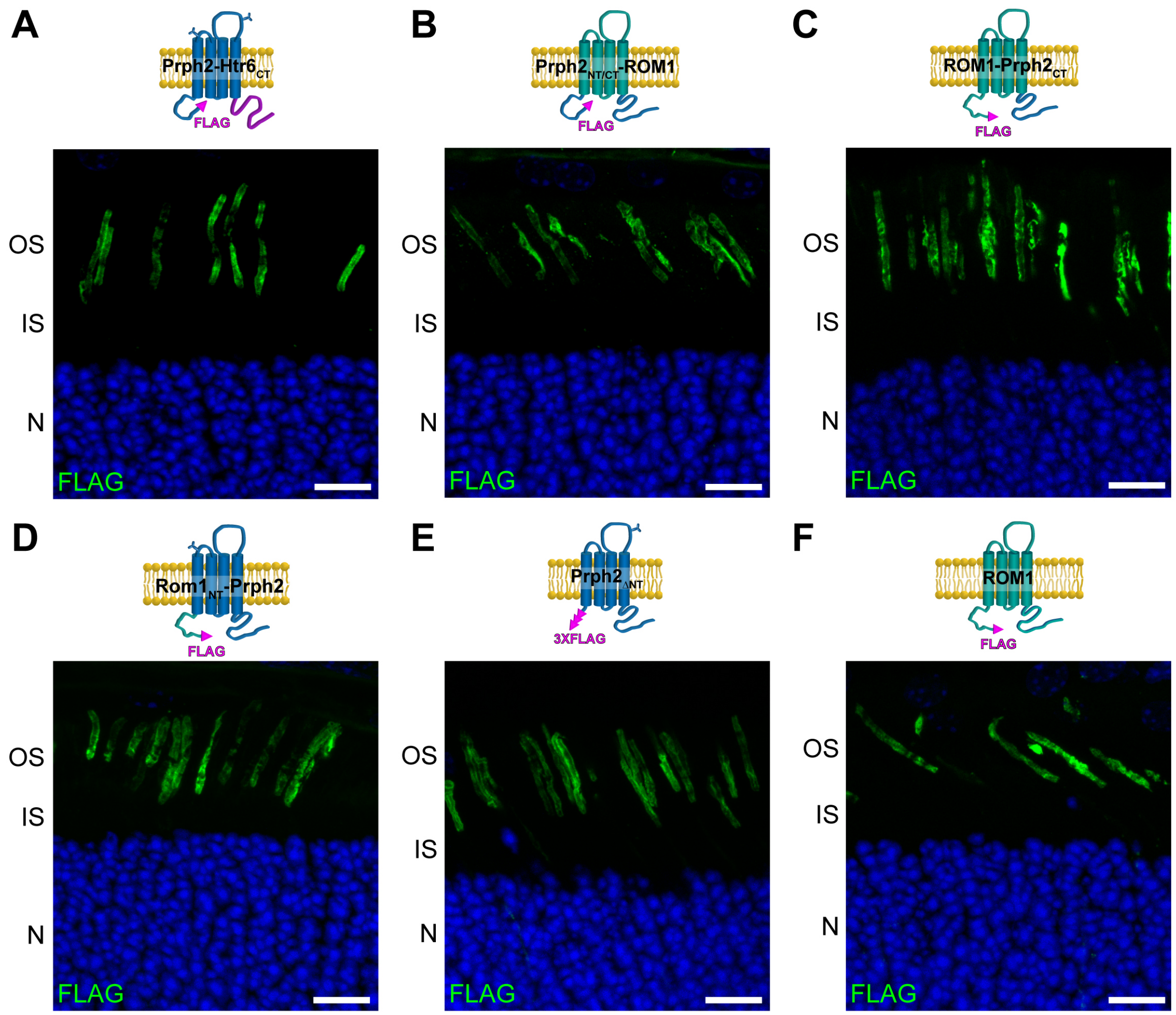


**Supplementary Figure 4.** WT mouse retinal cross-sections expressing **A.** Prph2-Htr6$_{CT}$, **B.** Prph2$_{NT/CT}$-ROM1, **C.** ROM1- Prph2$_{CT}$, **D.** ROM1$_{NT}$-Prph2, **E.** Prph2$_{\Delta NT}$ or **F.** ROM1. Retinal sections immunostained for FLAG. Nuclei counterstained in blue. Scale Bar, 10 μm.

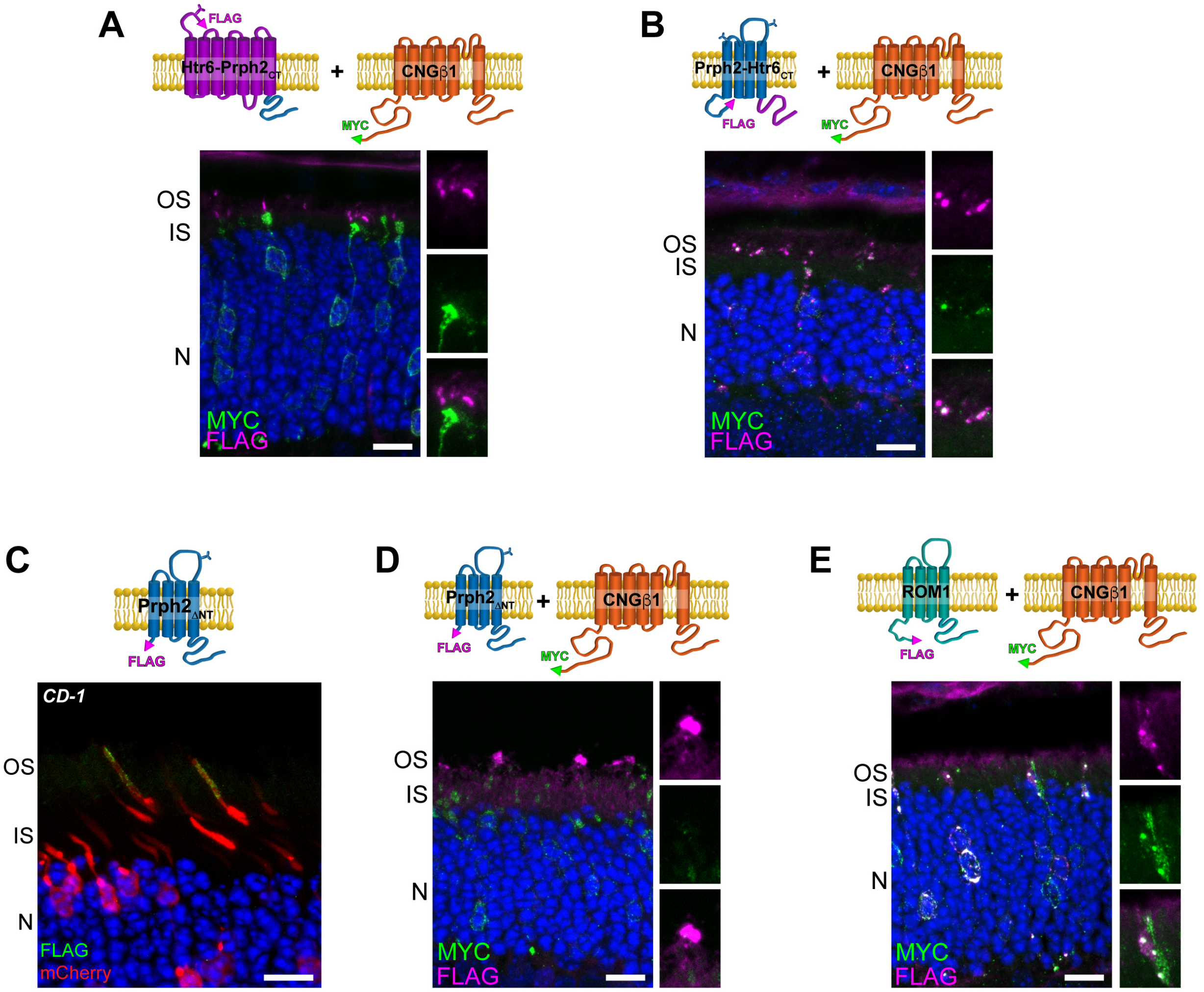


**Supplementary Figure 5.** *Rds*$^{-/-}$ mouse retinal cross-sections co-expressing MYC-CNGβ1 and **A.** Htr6-Prph2$_{CT}$ or **B.** Prph2-Htr6$_{CT}$. Retinal sections immunostained for FLAG and MYC. **C.** WT mouse retinal cross-sections expressing Prph2$_{\Delta NT}$ and mCherry. Sections immunostained for FLAG. **D.** WT mouse retinal cross-sections expressing Prph2$_{\Delta NT}$ and MYC-CNGβ1. Sections immunostained for FLAG and MYC. **E.** WT mouse retinal cross-sections expressing ROM1 and MYC-CNGβ1. Sections immunostained for FLAG and MYC. Nuclei counterstained in blue. Scale Bar, 10 μm.

### Figure 1C

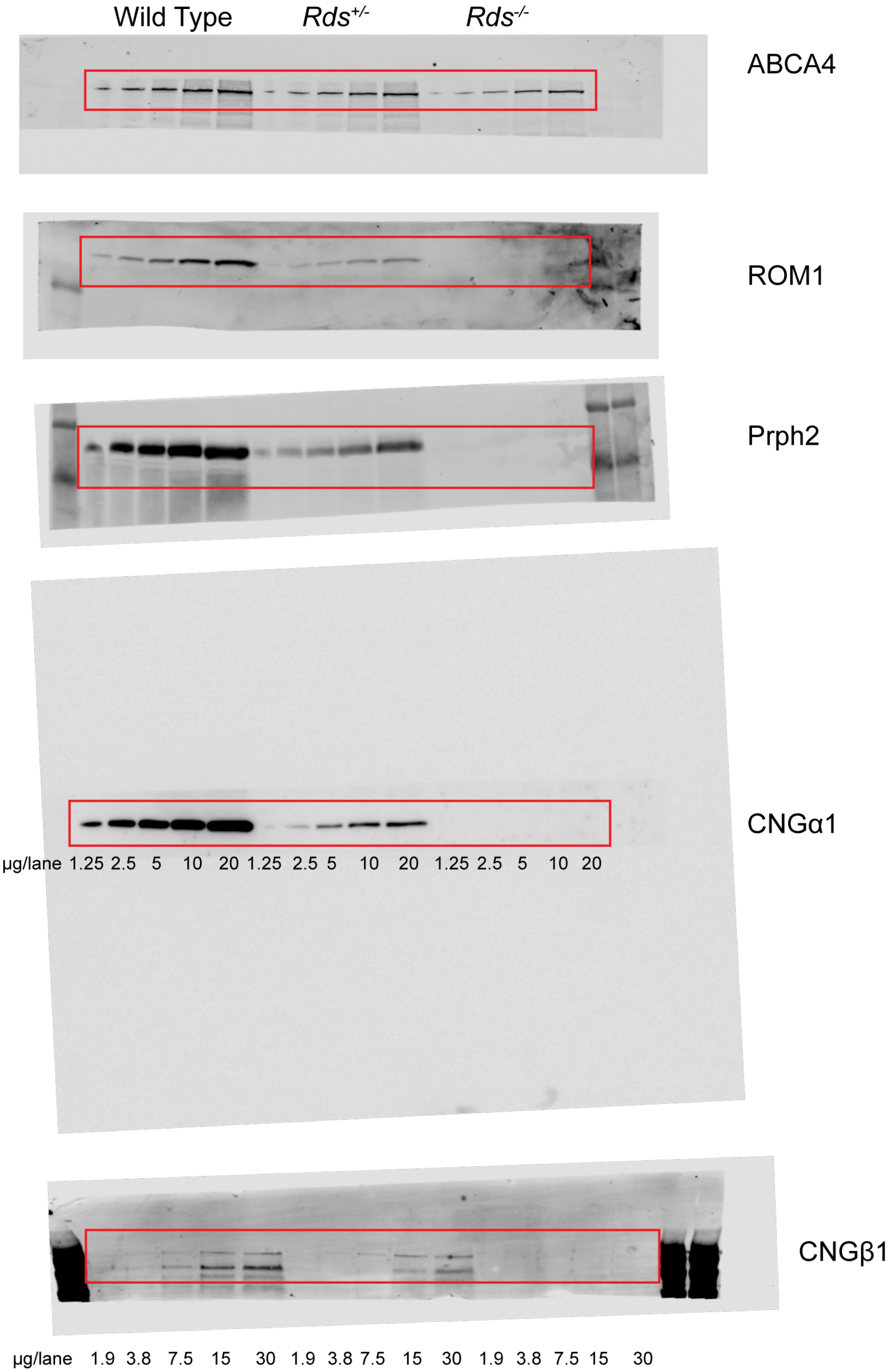
Wild Type
Rds+/-
Rds-/-
ABCA4
ROM1
Prph2
CNGα1
µg/lane 1.25 2.5 5 10 20 1.25 2.5 5 10 20 1.25 2.5 5 10 20
CNGβ1
µg/lane 1.9 3.8 7.5 15 30 1.9 3.8 7.5 15 30 1.9 3.8 7.5 15 30

**Figure 1E**

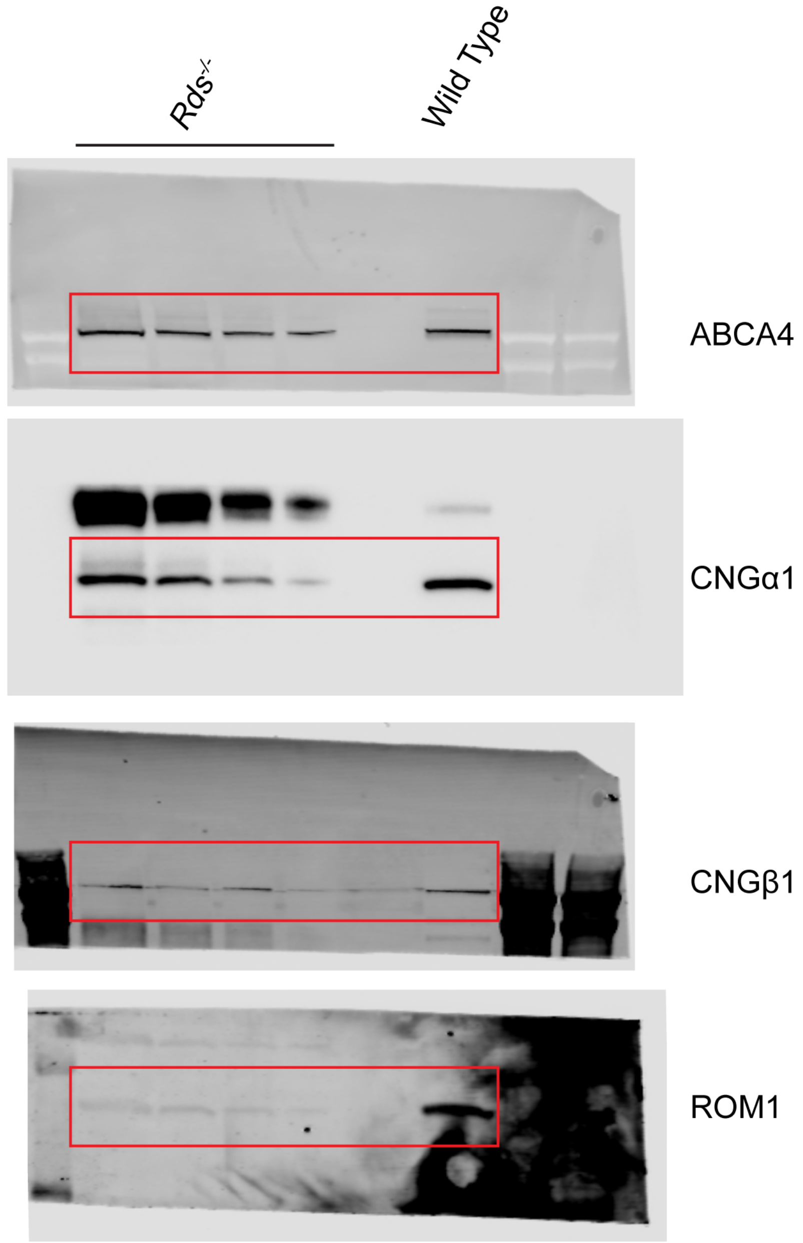

### Figure 3A

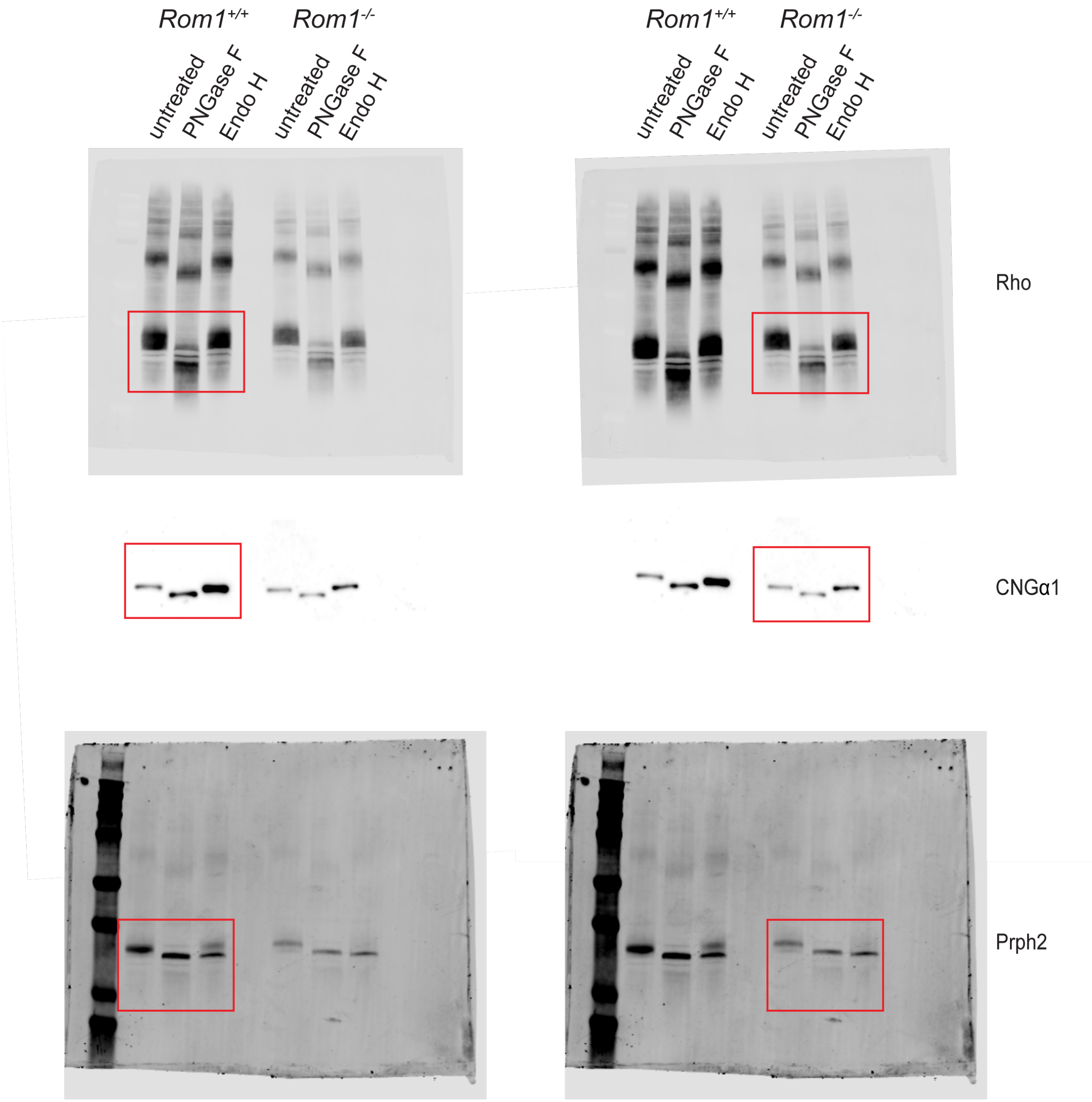


**Supplementary Figure 6.** Uncropped Western blot data

| CONSTRUCT | DESCRIPTION | PRIMER NAME | PRIMER SEQUENCE | FIGURE |
|---|---|---|---|---|
| MYC-CNGβ1 | Full-llength mouse CNGβ1 with C-terminal MYC tag | Cngb1-myc-R | CAGATCCTCTTCTGAGATGAGTTTTTGTTCCTCCGCCCCCTCTT | 2, 4, 5, Supp. Fig4, Supp. Fig5 |
| | | Not1-myc-R | GTGCGGCCGCCTACAGATCCTCTTCTGAGATGAGTTTTTGTTC | |
| $ROM1_{NT}$-Prph2 | FLAG-tagged ROM1 N-terminus (AA 1-21) fused to mouse peripherin-2 at TM domain 1 (AA 23-346) | Age1-Flag-Rom1-F | ATCCACCGGTAAGCATGGACTACAAGGACGACGACGACAAGGCGCCG | 4D, Supp. Fig3D |
| | | Rom1NT-Per-F | GTTTGGCACAGGGCATCTGGATGAACTGGCTGTCCGTG | |
| | | Rom1NT-Per-R | CACGGACAGCCAGTTCATCCAGATGCCCTGTGCCAAAC | |
| FLAG-$Prph2_{\Delta NT}$ | FLAG tagged peripherin-2 with N-terminus deleted up to TM1 (AA 18-346) | Flag-dNterm Prph2-F | GATCCACCGGTAAGCATGGACTACAAGGACGACGACGACAAGATGAACTGGCTGTCCGTG | Supp. Fig5 |
| 3xFlag-$Prph2_{\Delta NT}$ | 3xFLAG tagged peripherin-2 with N-terminus deleted up to TM1 (AA 18-346) | Age1-3XFLAG-F | GATTGCACCGGTAAGCATGGACTACAAAGACCATGACGGTGATTATAAAGATCATGACAT | 4E, Supp. Fig3E |
| | | 3XFLAG-dPerNT-F | TATAAAGATCATGACATCGACTACAAGGATGACGATGACAAGCAGGGGCTCTGGCTTATG | |
| Prph2-$Htr6_{CT}$ | FLAG-tagged peripherin (AA 1-290) fused to targeting region of Htr6 C-terminus (AA 392-440) | Per(T4)-Htr6CT-F | CGCTACCTCCACACAGCGCTGCTGCAGCTCACAGCCCAGC | Supp. Fig 2B. Supp. Fig3A |
| | | Per(T4)-Htr6CT-R | GCTGGGCTGTGAGCTGCAGCAGCGCTGTGTGGAGGTAGCG | |
| | | NotI-Htr6CT-R | GAGTGCGGCCGCTCAGTTCATGGGGGAACC | |
| ROM1-$Prph2_{CT}$ | FLAG-tagged mouse ROM1(AA 1-289) fused to mouse PerCT (AA 287-346) | Age1-Flag-Rom1-F | ATCCACCGGTAAGCATGGACTACAAGGACGACGACGACAAGGCGCCG | 6A, Supp. Fig3C |
| | | Rom1-PerCT-F | GCTCCTTGGTTTGCGGTATTTGCACACAGCGCTGGAGAGTGTG | |
| | | Rom1-PerCT-R | CACACTCTCCAGCGCTGTGTGCAAATACCGCAAACCAAGGAGC | |

**Supplementary Table 2.** Constructs and primer sequences.
